# Pathogen-dependent biocontrol activity of *Chlorella sorokoniana* aqueous extracts against fungal and oomycete plant pathogens

**DOI:** 10.64898/2026.08.05.742703

**Authors:** Ramona Mihaela Ciubotaru, Miguel Claro, Catarina Viana, Rui Figueiras, Pedro Rosa, Beatriz Duarte, Aghata C.R. Charnobay, Cláudia Rato, Sara Tedesco, Samira Andrade, Luísa Coelho, Florinda Gama, Mário Reis, Sandra Correia, Cristina Azevedo

**Author notes:** These authors contributed equally to this work (Miguel Claro, Catarina Viana). Corresponding author: C. Azevedo.

## Abstract

Fungal and oomycete plant pathogens are major drivers of yield losses worldwide and are spurring the search for sustainable alternatives to synthetic pesticides. Algae, in general, and microalgae, in particular, represent a promising source of bioactive compounds for crop protection. Despite this, their efficacy across different host–pathogen systems remains poorly characterised. This study evaluated the biocontrol potential of the aqueous extract of *Chlorella sorokoniana* against three economically important phytopathogens using complementary *in vitro*, *ex vivo*, and *in planta* assays. The strongest activity was observed against *Magnaporthe oryzae*, with the extract reducing fungal growth by approximately 70% *in vitro*, inhibiting appressorium formation by 55%, suppressing lesion development on detached rice leaves by more than 75%, and reducing rice blast severity by 64.5% as a preventive foliar treatment. In contrast, the extract showed little or no direct *in vitro* antifungal activity against *Pythium ultimum* and *Rhizoctonia solani*, yet it significantly reduced disease severity *in planta* by 13-37% and 48–55%, respectively. The contrasting responses among pathosystems suggest that *C. sorokoniana* aqueous extracts act through different mechanisms depending on the pathogen and the crop, combining direct antifungal activity against *M. oryzae* with plant-associated protective effects against soil-borne pathogens. These findings highlight the importance of evaluating candidate biocontrol products using complementary *in vitro* and *in planta* approaches, as laboratory antimicrobial assays alone may substantially underestimate their agricultural potential. The broad-spectrum protection achieved with an unrefined aqueous extract further supports *C. sorokoniana* as a promising source of sustainable crop protection products and provides a strong foundation for future mechanistic studies, formulation development, and field validation.

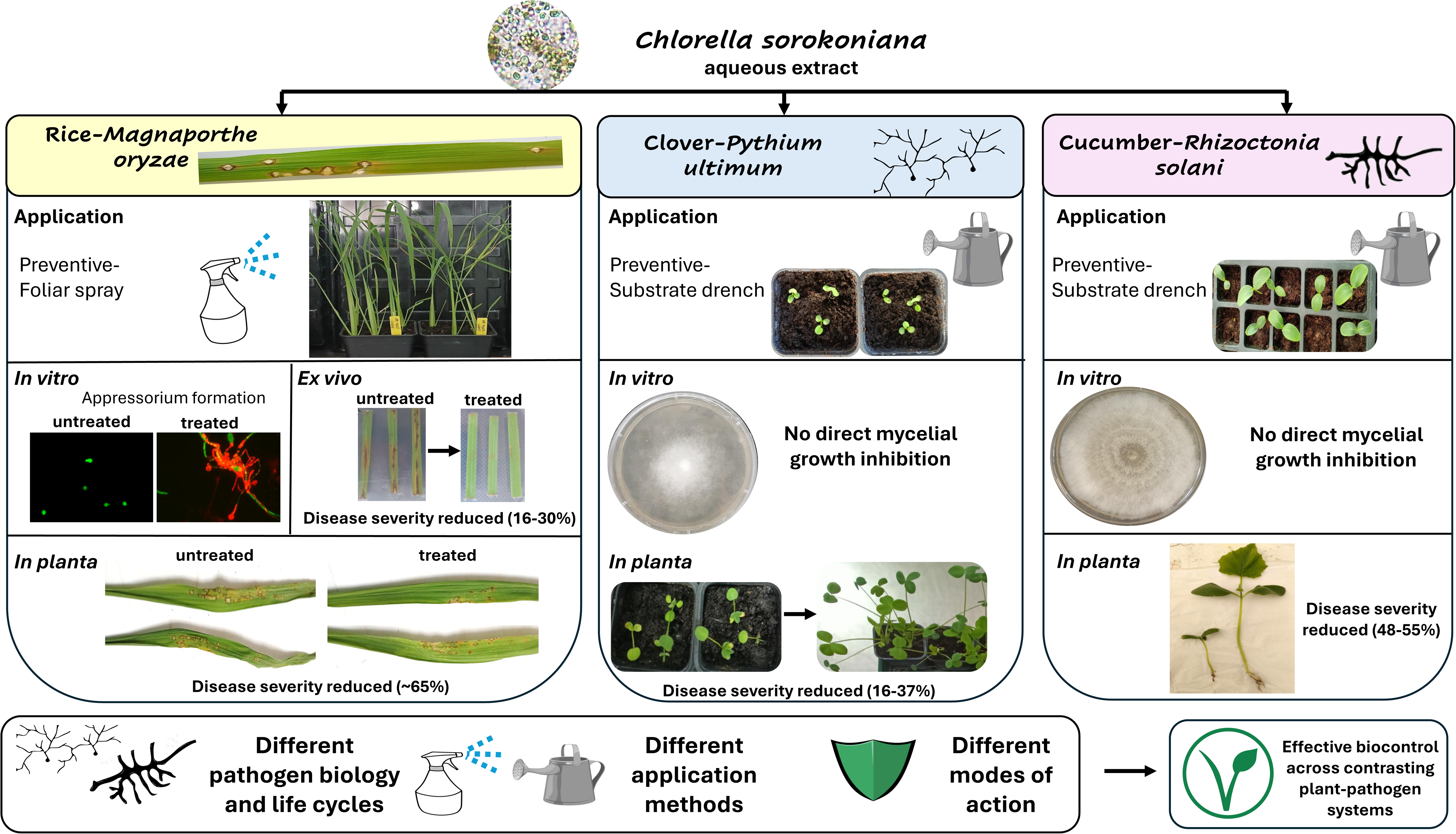

## 1. Introduction

Plant diseases caused by fungi and oomycetes limit agricultural yields, accounting for global yield losses of 20%-40% (Agrios, 2005; Savary et al., 2019). Among the most economically important phytopathogens are *Magnaporthe oryzae*, *Pythium ultimum*, and *Rhizoctonia solani* (Dean et al., 2012; Kamoun et al., 2014). The fungus *M. oryzae*, the causative agent of rice blast disease, is one of the most destructive fungal pathogens worldwide. Rice is the primary dietary staple for more than half of the world’s population; consequently, annual yield losses of 10–35% caused by the disease have a major impact on global food security (Asibi et al., 2019; Dean et al., 2012). *Pythium ultimum*, a soilborne oomycete, and *R. solani*, a soilborne fungal pathogen, are major causal agents of damping-off and root rot in numerous economically important crops (Arora et al., 2021; Kamoun et al., 2014; Matias et al., 2024). *Rhizoctonia solani* additionally causes sheath blight and infects more than 200 plant species, including rice, potato, and maize (Matias et al., 2024). The management of these diseases relies heavily on synthetic fungicides, whose intensive use has contributed to environmental contamination, biodiversity loss, and the emergence of resistant pathogen populations (Carvalho, 2017; Pathak et al., 2022). In response, the European Green Deal and the Farm to Fork Strategy aim to reduce pesticide use by 50% by 2030, highlighting the need to develop effective, sustainable crop protection alternatives (Jacquet et al., 2022).

Biological control has emerged as a sustainable alternative to synthetic pesticides. Although bacterial and fungal biocontrol agents are commercially available, their performance may vary under field conditions (Köhl et al., 2019, Palmieri et al., 2022). Seaweeds have been used in agriculture for centuries, primarily as organic fertilisers and soil amendments, owing to their ability to improve soil fertility, enhance nutrient availability, and promote plant growth (Khan et al., 2009). Microalgae have recently attracted attention as a source of bioactive metabolites with antimicrobial, plant-biostimulant, and defence-eliciting properties. Compounds such as polysaccharides, fatty acids, phenolics, peptides, and terpenoids have been associated with inhibition of pathogen development and activation of plant immune responses (Vehapi et al., 2020; Righini et al., 2022; Bhardwaj et al., 2025). In addition, microalgae can be produced sustainably using limited resources, increasing their attractiveness for crop protection applications (Spolaore et al., 2006; Colla & Rouphael, 2020). Among green microalgae, *Chlorella sorokoniana* has received particular attention for its rapid growth and rich composition of bioactive metabolites, including carotenoids, fatty acids, phenolic compounds, peptides, terpenoids, and alkaloids (Vehapi et al., 2020; Ouzakar et al., 2025). Previous studies have reported in *vitro* antifungal activity of *C. sorokoniana* against several phytopathogens, including *Fusarium oxysporum*, *Colletotrichum acutatum* and *Alternaria alternata* (Ferreira et al., 2021; Schmid et al., 2022; Ouzakar et al., 2025), while suppressive effects have also been demonstrated under *in planta* conditions for *F. oxysporum* (Viana et al., 2024). Despite these promising findings, most studies evaluated *C. sorokoniana* against pathogens exclusively *in vitro*, making it difficult to distinguish between direct antimicrobial activity and plant-mediated protection. Comparative studies integrating *in vitro*, *ex vivo*, and *in planta* assays across multiple pathosystems remain scarce, limiting our understanding of the mechanisms underlying disease suppression and the broader applicability of microalgae-based crop protection products.

This study investigated the biocontrol potential of aqueous extracts of *C. sorokoniana* using *M. oryzae*-rice as the primary pathosystem. A combination of *in vitro*, *ex vivo*, and *in planta* assays was used to determine whether direct antifungal activity translated into disease suppression and to elucidate the underlying mode of action. To assess the broader applicability of the algae extract, complementary *in vitro* and *in planta* assays were also performed against additional economically important pathogens, including *P. ultimum* and *R. solani*.

## 2. Materials and Methods

The experimental work was conducted within a collaborative framework between two research institutions, each contributing with complementary facilities, expertise, and pathosystem-specific trials. At InnovPlantProtect (38°53′03.4′′ N, 7°08′22.0′′ W), *in vitro, ex vivo* and *in planta* assays were performed in the *M. oryzae*–rice and *P. ultimum*–clover pathosystems, within the institution’s dedicated laboratories and greenhouse facilities. GreenCoLab carried out *in vitro* and *in planta* assays involving the *R. solani* pathosystem, with the *in planta* trials conducted in the greenhouse facilities of the Faculty of Sciences and Technology (FCT), Gambelas *Campus*, University of Algarve (37°02′35.45′′ N, 7°58′20.64′′ W). Together, this coordinated approach enabled the evaluation of *C. sorokoniana* extract treatments across three crop-pathogen systems, combining the experimental infrastructure and expertise of both institutions.

### 2.1 Microalgae biomass

*Chorella sorokoniana* dry biomass used in this study was supplied by Allmicroalgae Natural Products S.A. (Leiria, Portugal), batch L201910079 in accordance with the company’s established industrial protocols. After cultivation, the biomass was harvested by membrane filtration and spray-dried to obtain a fine powder, which was stored in sealed bags containing an oxygen absorber until use.

### 2.2 Preparation of *C. sorokoniana* aqueous extracts for *in vitro* assays

Aqueous extracts of *C. sorokoniana* biomass were prepared following an extraction workflow adapted from Schmid et al. (2022). Briefly, stock suspensions (10 g L⁻¹, w/v) were prepared by dispersing dried algae biomass in distilled or Milli-Q water and kept under continuous agitation at 350 rpm overnight in the dark. Depending on the experimental setup, extraction was performed either in Erlenmeyer flasks or centrifuge tubes. Following extraction, suspensions were subjected to two consecutive centrifugation steps (905–950 RCF) to remove particulate material. The recovered supernatants were clarified by sequential filtration through filters of progressively smaller pore sizes: 20, 6-10, and 4-7 µm (ᴓ150mm; LabKit Prat Dumas A015107, A015613, and A015110, respectively).

For the *in vitro* inhibition assays against *M. oryzae* and *P. ultimum,* the clarified aqueous algal extracts were sterilised under a laminar-flow hood by filtration through a sterile 0.2 µm membrane filter, then aliquoted into 96-well deep-well plates and stored at −20 °C until use*. For* the R*. solani* assay, the clarified extract was subjected to vacuum-assisted membrane filtration through a sterile 0.7 µm filter prior to final sterilisation using 0.2 µm bottle-top filters (Fisherbrand, England), aliquoted into 15 mL centrifuge tubes and stored as above.

### 2.3 *In vitro* pathogen growth inhibition assays

All growth inhibition assays across the different pathogens were performed with biomass derived from the same *C. sorokoniana* batch, and the experimental procedures were adapted according to the biological characteristics of each pathogen and assay platform. The main experimental conditions, treatment concentrations, incubation parameters, and evaluation criteria used in the *in vitro* assays are summarised in Table 1.

#### 2.3.1 Magnaporthe oryzae assays

Two *M. oryzae* Portuguese isolates were transformed to express the green fluorescent protein GFP, PR0009 (Gladieux et al., 2018) and M22.7 (Rosa et al, 2026). Briefly, PR0009-GFP was generated through *Agrobacterium tumefaciens*-mediated transformation (Rho et al., 2001) using the AGL-1 strain containing pCAMBgfp (Sesma and Osbourn, 2004), which expresses SGFP (Green fluorescence gene encoding a GFP variant that contains aserine-to-threonine substitution at amino acid 65). Stable, GFP-expressing clones were selected in the presence of 250 µg/mL hygromycin B, and fluorescence was observed under ultraviolet light. Independent transformants with indistinguishable morphology and very similar growth rates to their wild types (WTs) were selected (data not shown). The PR0009-GFP strain (Supplementary Figure 1) was used for all *in vitro* fluorescence and microscopy assays, whereas the wild-type strains Guy11 and M22.7 (Rosa et al., 2026) were used for detached-leaf and whole-plant infection assays. The isolates were routinely maintained on complete medium (CM; Talbot et al., 1993) or rice flour (RF; Le Naour-Vernet et al., 2023) agar (20 g/L organic rice flour, 2.5 g/L yeast extract, 1.5% agar) at 24–25 °C for seven to ten days under a constant light source consisting of 2 white and 2 purple fluorescent lamps with an intensity of 80-90 µmol.m^-2^.s^-1^ to promote vegetative growth and conidiation. Conidia were harvested from sporulating cultures using sterile distilled water containing 0.02% (v/v) Tween-20, filtered to remove mycelial fragments, and adjusted to the appropriate concentration for each assay (Supplementary Table 1).

##### 2.3.1.1 Fluorescence-based growth inhibition microplate assay

The antifungal activity of the *C. sorokoniana* aqueous extracts was assessed using a high-throughput GFP-based fluorescence assay, in which fluorescence served as a proxy for conidial germination and mycelial growth. Conidia from 7-day-old PR0009-GFP cultures were harvested in sterile distilled water containing 0.02% (v/v) Tween-20 and adjusted to 1 × 10³ conidia mL⁻¹. Assays were performed in black flat-bottom 96-well microplates (Thermo Scientific Nunc F96 Microwell) containing conidial suspensions and *C. sorokoniana* aqueous extracts at 0.1 and 1.5 g L⁻¹, in a final volume of 150 µL. Sterile distilled water served as the negative control, and non-inoculated wells were included for background correction. Each treatment consisted of four technical replicates. Green fluorescent protein (GFP) fluorescence (excitation 479 nm, emission 520 nm) was recorded every 30 min for 72–96 h at 25 °C using an Agilent BioTek Synergy H1 microplate reader under continuous orbital shaking (282 rpm).

##### 2.3.1.2 Fluorescence-based growth inhibition microplate assay

To determine whether *C. sorokoniana* aqueous extracts affected early infection-related development, conidial germination and appressorium formation were assessed and quantified by fluorescence microscopy using the PR0009-GFP strain. Conidial suspensions were prepared as described above and adjusted to 5 × 10⁴ conidia mL⁻¹. The suspensions were mixed with aqueous extracts of *C. sorokoniana* at final concentration of 0.2 g L⁻¹ in the presence of 100 µM 1,16-hexadecanediol, a cutin analogue that stimulates germination and synchronizes appressorial development *in vitro*. Control suspensions were prepared identically without the algae extract. Aliquots (15 µL) were placed on borosilicate coverslips (Epredia®, 20×20 mm #1) and incubated in sealed humid chambers for 24 h to evaluate early and mature appressorium formation, respectively. Prior to microscopy, samples were stained with propidium iodide (1 µg mL⁻¹) to assess membrane integrity. Images were acquired at 20× magnification using identical acquisition settings for all treatments. Two biological replicates were analysed per treatment, and at least 50 conidia were observed per replicate.

##### 2.3.1.3 Fluorescence-based growth inhibition microplate assay

The *in vitro* growth inhibition of *P. ultimum* and *R. solani* in the presence of the aqueous extract of *C. sorokoniana* was evaluated. *Pythium ultimum* isolate was obtained from the InnovPlantProtect mycological collection, whereas *R. solani* was obtained from the shared mycological collection of the Mediterranean Institute for Agriculture, Environment and Development (MED) and GreenCoLab. All isolates were routinely maintained on potato dextrose agar (PDA; Allonne, France) at 25 ± 2 °C in the dark for 7 days before use. Dual-plate assays were performed using a modified agar-based method adapted from Viana et al. (2024). *C. sorokoniana* aqueous extract was prepared as described above and tested at final concentrations of 0.1 and 1.5 g L⁻¹. Sterile distilled water served as the negative control.

For *P. ultimum*, the aqueous extract was incorporated at 1:1 (v/v) into molten double-strength PDA before plate pouring. Plates were inoculated with the pathogen no longer than 24 hours later, and kept at 4°C in the dark. For *R. solani*, 450 µL of extract was evenly spread over the surface of solidified 90 mm PDA plates using a sterile Drigalski spatula 24 h before inoculation with the pathogen. For plate inoculation, 5 or 6.5-mm mycelial plug from the actively growing colony margin of the pathogens *P. ultimum* and *R. solani*, respectively, were placed at the centre of each plate. Mycelium plugs were also grown on 1xPDA plates as a control. Plates were incubated under the pathogen-specific conditions as described in Supplementary Table 1. Colony growth was assessed by measuring colony diameter along two perpendicular axes, before the colonies in control plates reached the border, which normally occurred after 72 h. The final colony diameter grown upon algae treatment (T) was calculated as the mean of both measurements, and the percentage of growth inhibition relative to the control (C) was determined according to the following equation: Growth inhibition (%) =[(C-T)/C] x 100. Each treatment consisted of three biological replicates for *P. ultimum* and five for *R. solani*.

### 2.4 Treatment evaluation on rice detached leaves infected with *Magnaporthe oryzae*

To determine the efficacy of *C. sorokoniana* aqueous extract in controlling rice blast infection, detached-leaf assays were performed using rice (*Oryza sativa* L. cv. Presto), which is susceptible to the *M. oryzae* isolate M22.7 (Rosa et al., 2026). Rice plants were grown from seed in a commercial substrate (SIRO® Relva) under controlled-environment conditions (22–24 °C, approximately 80% relative humidity, 16 h light/8 h dark photoperiod) until they reached 3–4 weeks of age. Fully expanded leaves were cut, surface-sterilised with 70% ethanol, rinsed with sterile distilled water, and immersed in the *C. sorokoniana* aqueous treatment. Excess solution was removed using sterile filter paper before the leaves were placed on a Petri dish containing 0.6% agar (w/v) supplemented with 2 ppm 6-benzylaminopurine (6-BAP) to delay senescence. Five technical replicates were prepared for each treatment and incubated overnight in the dark. The *M. oryzae* conidial suspension inoculum (1 × 10⁵ conidia mL⁻¹) was prepared as described in section 2.4.1.1 in sterile distilled water supplemented with 0.02% (v/v) Tween-20 and 0.25% (w/v) gelatine. Leaves were infected the following day by applying a 4 μL droplet of an *M. oryzae* conidial suspension (1 × 10⁵ conidia mL⁻¹) to the centre of the leaf blade, avoiding any mechanical wounding. Following infection, the Petri dishes-containing the infected leaves were incubated overnight in the dark and subsequently maintained under a 16 h light/8 h dark photoperiod for up to 14 days. Disease development was assessed by measuring lesion area at 7 and 14 days post-inoculation (dpi) using ImageJ software (version 1.54d, Java 1.8.0_345(64-bit)).

### 2.5 Treatment evaluation on whole plants

To further corroborate the efficacy of *C. sorokoniana* aqueous extracts in controlling rice blast infection, whole-plant experiments were performed. The *C. sorokoniana* biomass evaluated was derived from the same production batch. Experimental procedures were adapted to the biological characteristics of each host–pathogen system, including the treatment application method, inoculation procedure, and disease assessment strategy. The principal experimental conditions for each pathosystem are summarised in Supplementary Table 2.

#### 2.5.1 Experimental design

The biocontrol efficacy of *C. sorokoniana* aqueous extract was evaluated in three representative host–pathogen systems differing in pathogen biology and infection strategies: *M. oryzae*–rice, *P. ultimum*–Persian clover, and *R. solani*–cucumber, under controlled greenhouse or growth-chamber conditions. All experiments included untreated, uninfected controls; infected controls; and pathogen-specific synthetic fungicide controls (when possible). All experiments followed a completely randomised design, with the number of biological replicates varying according to the requirements of each pathosystem. Experimental units (pots) were randomly distributed and periodically repositioned throughout the experimental period to minimise positional effects. Disease severity was assessed using pathosystem-specific ordinal scales. For the *M. oryzae*–rice pathosystem, disease severity was determined at the final assessment of 7 dpi, whereas for the *P. ultimum*–Persian clover and *R. solani*–cucumber pathosystems, disease progression was summarised as the Area Under the Disease Progress Curve (AUDPC). Treatment efficacy was calculated relative to the untreated infected control using the following equation: Treatment efficacy was calculated relative to the untreated infected control using the following equation: Treatment efficacy (%) = (Disease severity in control – Disease severity in treatment) / Disease severity in control × 100. The main experimental conditions adopted for each host–pathogen system is summarised in Supplementary Table 2.

#### 2.5.2 Preparation of *Chlorella sorokoniana* aqueous extracts for in planta assays

Stock suspensions of *C. sorokoniana* aqueous extracts for *in planta* assays were prepared by dispersing dried algae biomass in water and agitating overnight in the dark at 350 rpm. Depending on the experimental setup, extraction was performed either in Erlenmeyer flasks or centrifuge tubes. The extract obtained is similar to ALLFERTIS, commercialised by Allmicroalgae as an organic biofertilizer https://www.allmicroalgae.com/en/biostimulants-microalgae-allfertis/). Treatment concentrations were chosen independently for each pathosystem, based on the objectives of the respective experiments and the protocols established for each host–pathogen system. For the clover-*P. ultimum* pathosystem, stock suspensions were prepared at 30 g L⁻¹ and diluted in tap water to a final working concentration of 2 g L⁻¹ to maximise the plant-mediated protective effects, despite the absence of direct antifungal activity *in vitro*. For the rice-*M. oryzae,* stock suspensions were prepared at 10 g L⁻¹, and the working concentration was selected based on the effective dose identified *in vitro*, with a slight increase for plant-associated assays (0.2 g L⁻¹). For the cucumber-*R. solani* pathosystem, stock suspensions were prepared at 10 g L⁻¹ and diluted to final working concentrations of 1.5 g L⁻¹.

#### 2.5.3 Treatment evaluation on whole rice plants / *Magnaporthe oryzae* pathosystem

The preventative efficacy of *C. sorokoniana* aqueous extract against rice blast was evaluated using rice (*Oryza sativa* L. cv. Bomba) grown in 1 L pots containing commercial substrate (SIRO® Relva) under controlled growth chamber conditions (22–24 °C, approximately 80% relative humidity, 16 h light/8 h dark photoperiod). Treatments were applied preventively, 24 h before pathogen inoculation, as a foliar spray at the vegetative stage (2–3 weeks after sowing). Approximately 4 mL of aqueous *C. sorokoniana* extract was applied per pot using an airbrush sprayer (Timbertech® ABPST05) fitted with a 0.5 mm nozzle. Each treatment consisted of three biological replicates, with one pot containing six rice plants considered as a single experimental unit. Non-infected and infected control plants were sprayed with sterile distilled water. For the infection, *M. oryzae* Guy11 (Leung et al, 1988) conidial suspension was prepared at 1 × 10⁵ conidia mL⁻¹ in sterile distilled water supplemented with 0.02% (v/v) Tween-20 and 0.25% (w/v) gelatine. Twenty-four hours after treatment application, plants were spray-infected with approximately 2 mL of conidial suspension per pot under the same spraying conditions. Infected plants were incubated at 25 °C under saturated humidity (dew) in darkness for 24 h and transferred to a growth chamber maintained at 25 °C, approximately 95% relative humidity, and a 16 h light/8 h dark photoperiod for 6–7 days to allow symptom development. Disease symptoms were evaluated 7 days after infection using a modified disease assessment framework adapted from Hensawang et al. (2017) and IRRI (2002). Unlike the original disease assessment, which combines lesion morphology and disease severity into a single ordinal scale, the framework separates these components into two independent assessment scales, each applied to every evaluated leaf. Scale 1 – Lesion morphology classified lesion development according to the diameter (⌀) of the spot using six ordinal classes: 0 (no lesions); 1 (0-1 mm); 3 (1–2 mm); 5 (2–4 mm); 7 (4–7 mm); and 9 (>7 mm). Scale 2 – Disease severity was classified by the percentage of leaf area exhibiting blast symptoms into six ordinal classes: 0 (0%); 1 (1–5%); 3 (5–25%); 5 (25–50%); 7 (50–75%); and 9 (75-100%). Representative photos were used to standardise scoring for both scales (Supplementary Figure 2). The dual-scale framework enables quantification of disease progression using complementary descriptors of lesion morphology and disease severity, allowing disease index calculations based on the ordinal scores of both parameters. For quantitative statistical analyses, only the disease severity scale was used because its classes represent predefined disease severity intervals. Following Del Ponte (2023), Willocquet (2023), and Chiang (2022), the ordinal disease severity scores were converted to the midpoints of their corresponding percentage intervals prior to statistical analysis. The converted values were then averaged across all assessed leaves to obtain a disease severity value for each plant. Plant-level values were subsequently used to calculate treatment means and standard deviations for statistical comparisons and graphical representation. This framework therefore extends conventional rice blast assessment by enabling both disease index calculations from complementary symptom descriptors and statistically robust disease severity analyses based on interval-derived percentage leaf area values.

#### 2.5.4 Treatment evaluation on whole Persian clover / *P. ultimum* pathosystem

The efficacy of *C. sorokoniana* aqueous extract against *P. ultimum* was evaluated using Persian clover (*Trifolium resupinatum* cv. Resal) grown in a controlled growth chamber. Seeds were pre-germinated on sterile filter paper moistened with distilled water for 24 h in the dark at room temperature (23 °C), then incubated under a 16 h light/8 h dark photoperiod for an additional 3 days.

*Pythium ultimum* inoculum was produced on sterilised organic millet. Hydrated millet grains were autoclaved, inoculated with six agar plugs (4 mm) from actively growing cultures, and incubated at 25 °C in darkness for approximately two weeks with periodic agitation to promote uniform colonisation. Commercial substrate (SIRO® Relva) was pasteurised at 83 °C for 30 min. Half of the pasteurised substrate was placed into 6 × 6 cm pots to form the bottom layer. The remaining substrate was thoroughly mixed with *P. ultimum*-colonised millet at 1% (w/w) and added as the upper layer to complete the pots. The *C. sorokoniana* aqueous extract was applied preventively by substrate drench (30 mL per pot) immediately before transplanting three pre-germinated Persian clover seedlings into each pot. Each treatment consisted of five biological replicates. Infected and non-infected controls received an equivalent volume of water, while the commercial fungicide Rival® (propamocarb as active ingredient) was included as a positive control and applied at 0.6 mL L⁻¹. Plants were maintained in a growth chamber under controlled conditions (14 °C day/9 °C night, approximately 95% relative humidity, 16 h light/8 h dark photoperiod) for one week before being transferred to a walk-in growth chamber (approximately 21 °C and 70% relative humidity) for the remaining two weeks of the experiment.

A disease severity scale adapted from You et al. (2017), itself based on the root disease severity scale developed by Wong et al. (1984), was further developed to assess damping-off caused by *P. ultimum* in Persian clover over a three-week period. The original protocol was extended from a single-end-point assessment of root disease to a physiological stage-specific disease severity scale that followed the progression of damping-off throughout plant development. This modified approach combined consecutive, non-destructive assessment of above-ground symptoms with a final destructive evaluation of root necrosis, using a four-point ordinal scoring system. During the first week after inoculation, disease severity was assessed using seedling survival, with 1 = alive and 4 = dead. During the second week, disease severity was evaluated based on two above-ground symptoms: plant height and leaf chlorosis. Plant height was scored as: 1-comparable to the non-inoculated control; 2-intermediate reduction in height; 3-severely stunted; and 4-dead plant. Height categories were established independently for each experiment after assessing the height of all plants relative to both control groups: not infected and infected. Plants with heights comparable to the non-infected control were assigned a score of 1, and clearly stunted plants, comparable to the infected control, were assigned a score of 3. The tallest plant within the stunted group defined the upper threshold for score 3, while the shortest plant within the healthy-height group defined the lower threshold for score 1. Plants with heights falling between these two thresholds were classified as score 2. Leaf chlorosis was scored as: 1-Healthy green plant comparable to the non-inoculated control; 2-≥50% green tissue remaining; 3-<50% green tissue remaining; 4-dead plant. During the third week, the same above-ground symptoms were reassessed, and root necrosis was evaluated after uprooting the plants. Root symptoms were scored as: 1-healthy root system comparable to the non-inoculated control; 2-≤ 50% root necrosis or reduction in root length; 3-> 50% root necrosis or reduction in root length; and 4-dead plant (Supplementary Figure 3). The disease severity scores obtained at each assessment were used to calculate the area under the disease progress curve (AUDPC) for each biological replicate using the standard trapezoidal method (Campbell and Madden, 1990), as implemented in the audpc() function of the agricolae package in R (de Mendiburu, F. 2023). Disease assessments corresponded to the first-week survival score, the mean disease severity score obtained from plant height and chlorosis during the second week, and the mean disease severity score obtained from plant height, chlorosis, and root necrosis during the third week.

#### 2.5.5 Treatment evaluation on whole cucumber / *Rhizoctonia solani* pathosystem

The efficacy of *C. sorokoniana* aqueous extracts against *R. solani* was evaluated in cucumber (*Cucumis sativus* cv. Marketmore) grown under greenhouse conditions with a natural photoperiod (approximately 11 h light/13 h dark), with micro-sprinkler irrigation to maintain substrate moisture. Peat substrate (Du Vitor®) was amended with calcium carbonate (4.5 g L⁻¹) to adjust the pH to approximately 6.5, following Viana et al. (2024), and distributed into 70 mL pots. *Rhizoctonia solani* was cultured on PDA for 7 days before inoculation. Each pot was infected by placing a 6.5 mm mycelial plug at the centre of the moistened substrate and incubating for 7 days before sowing to allow pathogen establishment. Treatments were applied preventively immediately after sowing by substrate drench with 7 ml of aqueous *C. sorokoniana* extracts at 1.5 g L⁻¹. Each treatment comprised twenty technical replicates, with one pot containing one plant considered as a single experimental unit. Infected and uninfected controls received only water, while the commercial synthetic fungicide Moncut SC® (Massó®) served as the positive control and was applied at 0.75 L ha^-1^ according to the manufacturer’s recommendations. Plants were maintained under greenhouse conditions throughout the experiment. Disease development was assessed weekly according to the disease assessment protocol of Matias et al. (2024). Germination, seedling survival, plant growth, chlorosis, and plant mortality were monitored throughout the experiment. At the final assessment, non-germinated seeds were recovered to confirm pre-emergence damping-off caused by *R. solani*. Disease severity was scored using a five-class ordinal scale: 1-No visible symptoms; 2-Minor symptoms; 3-Severe symptoms; 4 – Post-emergence damping-off or plant death; and 5-Pre-emergence damping-off. Disease severity scores were subsequently used to calculate the AUDPC.

## 3. Statistical analysis

All statistical analyses were performed in R (version 4.3.3). Data from independent experiments were pooled prior to statistical analysis for all *in vitro*, *ex vivo* and *in planta M. oryzae* assays, whereas independent experiments for the *P. ultimum* and *R. solani* pathosystems were analysed separately. Results are presented as mean ± standard deviation (SD). Differences were considered statistically significant at *p* < 0.05.

### 3.1 *In vitro* assays

For the fluorescence-based microplate growth inhibition assay, blank-corrected fluorescence data were analysed by one-way ANOVA, and inhibition at each treatment concentration was compared with the theoretical value of 0 using a two-sided one-sample Student’s t-test. Conidial germination and appressorium formation data were analysed using generalised linear models (GLMs) with a binomial error distribution and logit link function. For dual-plate growth inhibition assays, treatment effects were analysed using one-way analysis of variance (ANOVA) followed by Dunnett’s multiple comparisons test against the untreated control.

### 3.2 Detached-rice leaves treatment assays

For detached leaf assays, treatment effects on lesion inhibition were assessed using a two-sided one-sample Student’s t-test against the theoretical value of 0.

### 3.3 *In planta* treatment assays

Disease symptoms in the *M. oryzae*–rice pathosystem were independently scored using two ordinal scales describing lesion morphology (spot diameter) and disease severity (percentage of diseased leaf area). The dual-scale framework enables both disease index calculations and quantitative assessment of disease severity. For statistical analyses, only the disease severity scale was used because its classes represent predefined severity intervals. Following Willocquet et al. (2023), ordinal scores were converted to the midpoint of their corresponding percentage intervals before analysis. Data were assessed for normality (Shapiro–Wilk test) and homogeneity of variance (Levene’s test). When assumptions were met, treatment effects were analysed by one-way ANOVA followed by Tukey’s HSD test. Otherwise, Welch’s ANOVA or the Kruskal–Wallis test was applied, with Dunn’s post hoc test where appropriate. Count data were analysed using generalised linear models (GLMs) with a quasi-Poisson distribution, and differences in proportions were assessed using Fisher’s exact test. Data were assessed for normality (Shapiro– Wilk test) and homogeneity of variance (Levene’s test). When assumptions were met, treatment effects were analysed by one-way ANOVA followed by Tukey’s HSD test. Otherwise, Welch’s ANOVA or the Kruskal–Wallis test was applied, with Dunn’s post hoc test where appropriate. For the *P. ultimum*–clover and *R. solani*–cucumber disease progression, the area under the disease progress curve (AUDPC) was quantified, and treatment effects were analysed by one-way ANOVA followed by Tukey’s HSD test.

## 4. Results

### 4.1 Inhibition of fungal growth by *Chlorella sorokoniana* aqueous extract *in vitro*

#### 4.1.1 Magnaporthe oryzae

We investigated the effect of *C. sorokoniana* aqueous extracts on inhibiting *M. oryzae* growth *in vitro* using a fluorescence-based 96-well microtiter plate assay with the GFP-expressing strain PR0009-GFP. In this assay, an increase in fluorescence over time reflects spore germination and hyphal development. Treatment with *C. sorokoniana* aqueous extract at 0.1 g L⁻¹ significantly reduced fungal biomass accumulation by approximately 70% relative to the untreated control (p < 0.001). Whereas the treatment at 1.5 g L⁻¹ resulted in approximately 25% inhibition (p < 0.001; Figure 1). As the lower concentration produced the strongest inhibitory effect, subsequent mechanistic assays focused on this concentration range.

**Figure 1.**
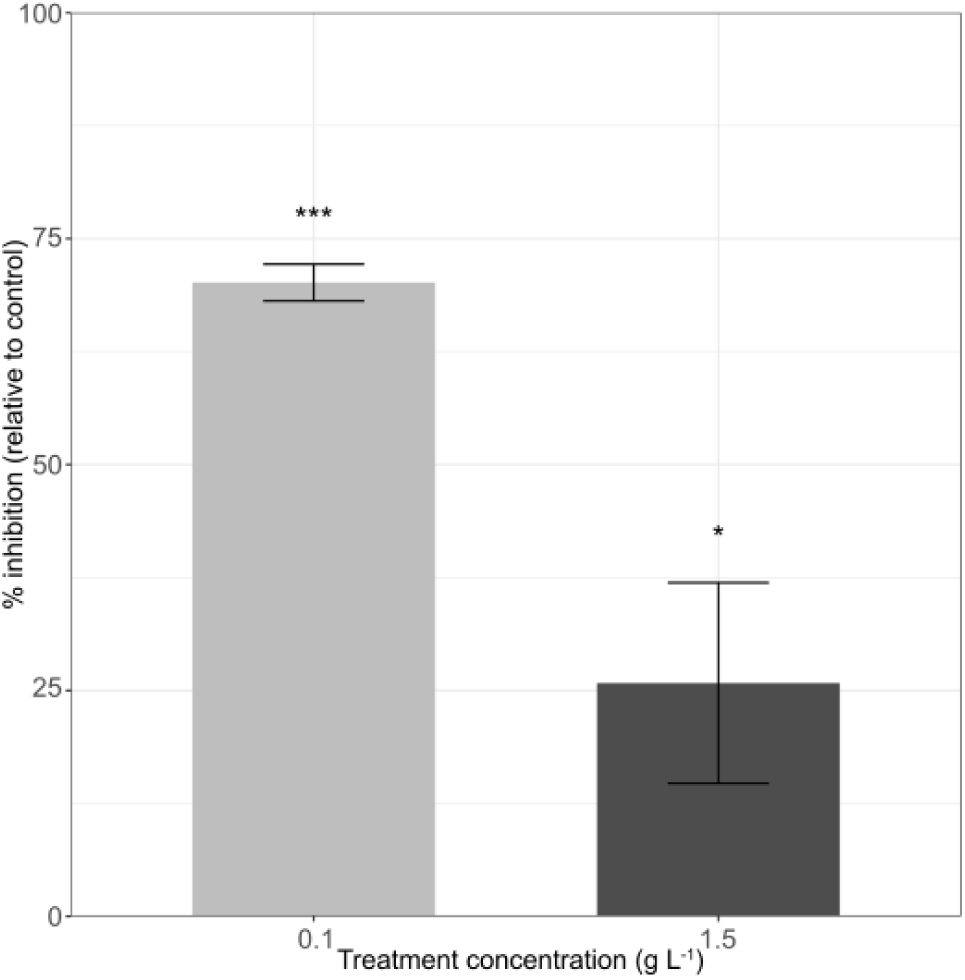
– High-throughput fluorescence-based *Magnaporthe oryzae* growth inhibition microplate assay to quantify the *in vitro* effect of *Chlorella sorokoniana* aqueous extract on *M. oryzae* growth inhibition (%). Comparison is made relative to the untreated control at 0.1 and 1.5 g L⁻¹. Data represented as mean ± SD correspond to pooled results from two independent experiments (three replicates per experiment). Statistical analyses were performed comparing each treatment with the untreated control using one-way ANOVA, followed by Dunnett’s multiple comparisons test. Asterisks indicate significant differences from the untreated control (* *p* < 0.05; *** *p* < 0.001).

To determine whether the extract interfered with the initial events required for host infection, we next investigated its impact on conidial germination and appressorium formation. Untreated conidia germinated and developed healthy appressoria with short germination tubes. However, conidia treated with the *C. sorokoniana* aqueous extract at 0.2 g L⁻¹ showed abnormal fungal development such as elongated germ tubes and deformities in the germination tubes/appressoria (indicated by the arrows; Figure 2A). Moreover, propidium iodide staining confirmed the death of *M. oryzae* cells and the loss of appressorium viability, a critical structure for host infection. To evaluate appressorium formation, *M. oryzae* spores were treated with 0.2 g L⁻¹ of *C. sorokoniana* aqueous extract (Figure 2B). The treatment significantly reduced appressorium formation by approximately 55% compared with the untreated control (*p* < 0.001, binomial GLM; Figure 2C). Although variability among biological replicates was observed, the reduction in appressorium formation was consistent across independent experiments.

**Figure 2.**
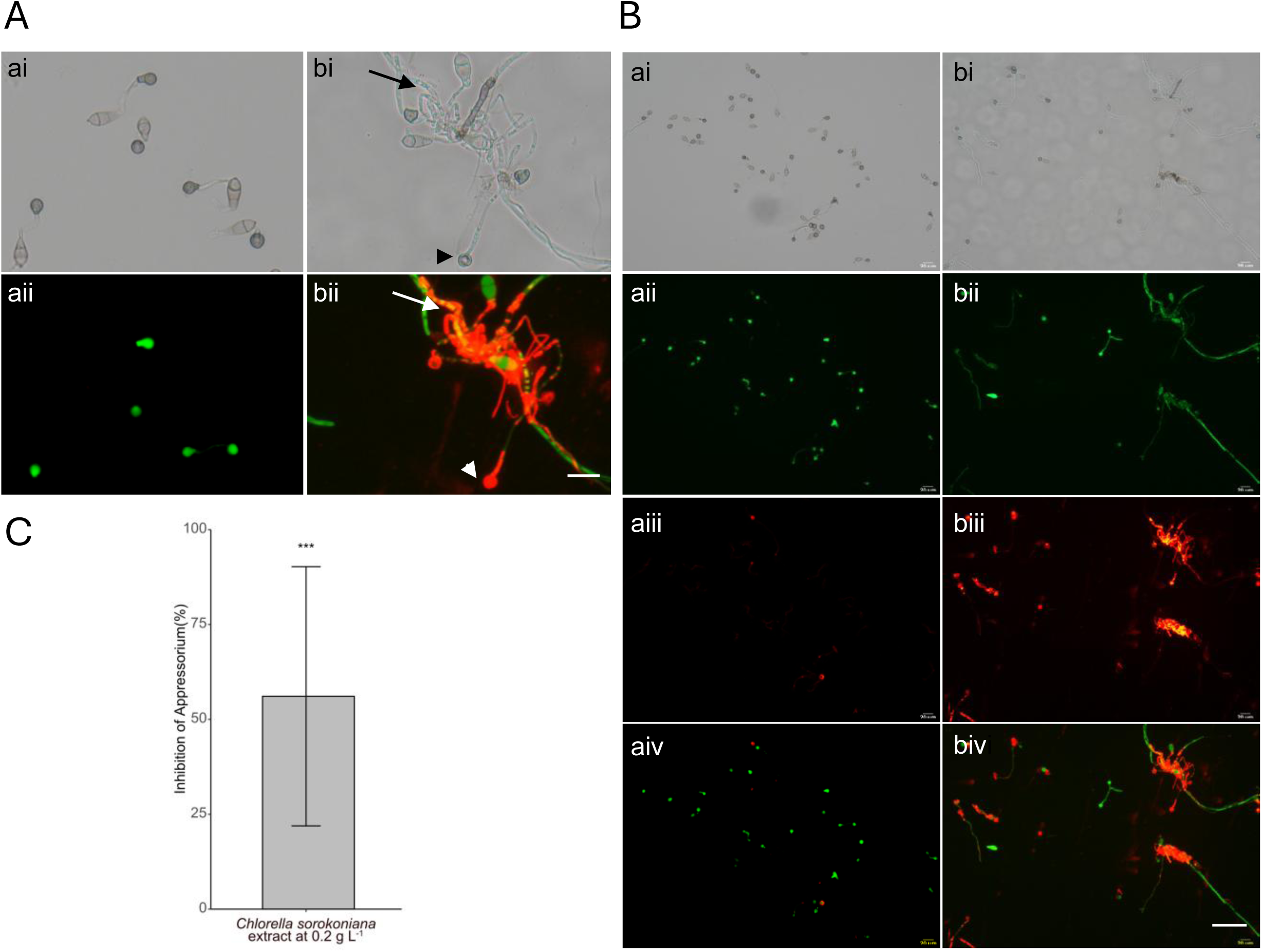
– *In vitro* assessment of the effect of *Chlorella sorokoniana* aqueous extract on *Magnaporthe oryzae* (PR9-GFP) spore germination and appressorium formation. In panels A and B, “a” represent the untreated conidia control, and “b” the conidia treated with *C. sorokoniana* aqueous extract (0.2 g L⁻¹). The spore solutions have been stained with propidium iodide to observe cell death (red channel). A) Effect of extract on *M. oryzae* conidia germination on a hydrophobic surface (arrows indicate deformities in germ tubes and triangles indicate deformities in appressoria, and both arrows and triangles indicate cell death). B) *M. oryzae* appressorium formation in brightfield (ai and bi) and darkfield (all other panels) on different channels: Green (aii and bii), Red (aiii and biii), and merged Green and Red channels (aiv and biv). C) Quantification of the percentage of inhibition of appressorium formation following treatment. Data represent results of two independent pooled experiments (50 spores counted per experiment; n = 2) and are presented as the mean ± SD. Statistical comparisons were performed against the untreated control using GLM with a binomial error distribution and logit link function (\*\*\**p* < 0.001). Scale 20 μm.

#### 4.1.2 Pythium ultimum

Oomycetes differ substantially from true fungi in their physiology and cell wall composition, and for this reason, *P. ultimum* was selected to evaluate whether the inhibitory activity of the extracts extended to this important group of plant pathogens. The antifungal activity of the *C. sorokoniana* aqueous extract against *P. ultimum* was first evaluated using a dual-plate assay. The extract showed negligible inhibition of mycelial growth at both tested concentrations (0.1 and 1.5 g L⁻¹), with mean inhibition values below 10% indicating little or no direct inhibitory activity (Supplementary Figure 4A).

#### 4.1.3 Rhizoctonia solani

To further assess the extract’s activity spectrum, its effects were evaluated against *R. solani*, a broad-host-range soilborne pathogen that causes severe diseases in many economically important crops. This pathogen was selected to determine whether the inhibitory activity extended to fungal pathogens with distinct lifestyles and infection strategies. A dual-plate assay showed that the *C. sorokoniana* aqueous extract exhibited negligible direct antifungal activity against *R. solani*. Mycelial growth inhibition remained below 4% at both tested concentrations (0.1 and 1.5 g L⁻¹), indicating no meaningful inhibition of fungal growth under *in vitro* conditions (Supplementary Figure 4B).

### 4.2 Inhibition of fungal growth by *Chlorella sorokoniana* aqueous extract *in planta*

The extract was subsequently evaluated *in planta* to determine whether its biological activity translated into disease suppression under plant conditions. As the extract inhibited the growth of *M. oryzae* but showed no direct activity against *P. ultimum or R. solani in vitro*, *in planta* assays were designed to address two objectives. First, the rice–*M. oryzae pathosystem* was used to confirm that the *in vitro* antifungal activity conferred protection against rice blast. Second, the Persian clover–*P. ultimum* and cucumber–*R. solani* pathosystems were included to investigate whether the extract could confer disease protection in the absence of direct antimicrobial activity, potentially through the activation of plant defence responses or other indirect plant-mediated mechanisms.

#### 4.2.1 Rice*-Magnaporthe oryzae* pathosystem

Given the extract’s promising antifungal activity against *M. oryzae in vitro*, its efficacy was assessed in rice using a stepwise approach. Initial experiments were performed on detached rice leaves to determine whether the extract could suppress disease development under controlled detached leaf conditions. Leaves treated with 0.2 g L⁻¹ extract before inoculation with *M. oryzae* developed significantly smaller lesions than the untreated infected controls (Figure 3A). Lesion development was reduced by more than 75%, corresponding to a significant inhibition of disease development (*p* < 0.001, Student’s T-test; Figure 3B).

**Figure 3.**
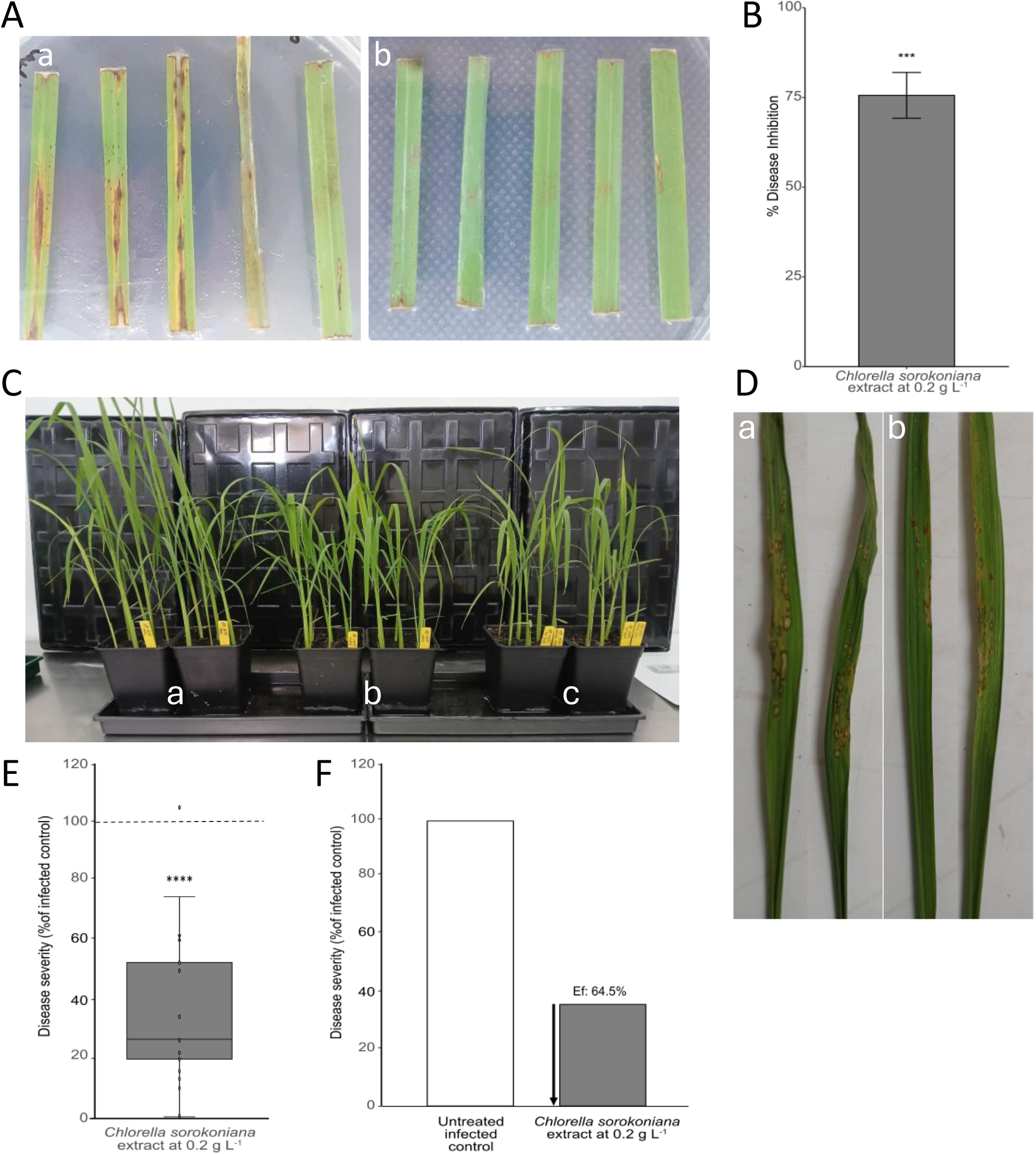
– Preventive activity of *Chlorella sorokoniana* aqueous extract against rice blast caused by *Magnaporthe oryzae*. A) Representative images of rice detached leaves corresponding to (a) untreated infected control and (b) *C. sorokoniana* extract (0.2 g L⁻¹)-treated infected. B) Quantification of the disease inhibition in rice detached leaf assays, expressed relative to the untreated infected control. C) Representative rice plants at 7 days post-inoculation (dpi): (a) non-inoculated untreated control, (b) untreated infected control, and (c) plants treated with *C. sorokoniana* extract 24 h before inoculation with *M. oryzae*. D) Representative images of individual rice leaves from the experiment in panel C corresponding to (a) untreated infected control and (b) *C. sorokoniana* extract (0.2 g L⁻¹)-treated infected. E) Disease severity in rice whole plants following foliar application of the extract 24 h before inoculation with *M. oryzae*, expressed relative to the untreated infected control (100%). Box-and-whisker plots summarise the biological replicates; the dashed line denotes the untreated infected control (100%), and the × indicates the mean. F) Quantification of the treatment efficacy (Ef), expressed as the percentage reduction in disease severity relative to the untreated infected control. Data in B, E and F represent the pooled results of two independent experiments (three biological replicates per experiment; n = 6) and are presented as the mean ± SD. Statistical comparisons were performed against the untreated infected control using Student’s t-test. Asterisks indicate significant differences from the untreated infected control (*** p < 0.001; **** p < 0.0001).

To evaluate whether the protective effects observed *ex vivo* translated to whole-plant responses, and therefore more closely resembled natural infection, preventive foliar applications of *C. sorokoniana* aqueous extract were assessed in the *M. oryzae*–rice pathosystem (Figure 3C-F). Preventive application of the extract at 0.2 g L⁻¹ significantly reduced rice blast severity reflected in the overall appearance of the plants at the final assessment (7 dpi), where infected, untreated plants exhibited less vigorous growth than both uninfected controls and infected, *C. sorokoniana*-treated plants (Figure 3C) and as can be observed in detail (Figure 3D).

To enable robust quantification of these visual differences, the modified disease assessment framework was validated by evaluating the relationship between lesion morphology (spot diameter) and disease severity (percentage of diseased leaf area) across five independent *M. oryzae*–rice experiments. The two assessment parameters were consistently and strongly correlated (Spearman’s ρ = 0.939–0.995, *p* < 0.001 across all experiments), indicating that both parameters captured the same progression of rice blast symptoms. These results validated the use of the percentage-based disease severity scale as the primary parameter for quantitative analyses while retaining the dual-scale framework for disease index calculations and comprehensive disease characterisation.

Using this validated disease severity scale, quantitative analysis confirmed that the preventive treatment significantly reduced rice blast severity when compared with the infected untreated control (*p* < 0.0001, Student’s t-test; Figure 3E). Disease severity was reduced to approximately 30% of the infected control, corresponding to a treatment efficacy of approximately 70% (Figure 3F). Lesion development was consistently reduced across biological replicates, demonstrating that the inhibitory effects observed on fungal growth and appressorium formation *in vitro* translated into substantial suppression of rice blast in the whole plant.

#### 4.2.2 Persian clover-*Pythium ultimum* pathosystem

Next, we investigated whether the extract could also exert plant-mediated effects, thereby contributing to disease suppression independently of direct antimicrobial activity. As the extract showed no direct inhibitory activity against *P. ultimum in vitro*, the *P. ultimum*–Persian clover pathosystem provided an ideal model to evaluate plant-mediated effects. Representative images of the assay are shown in Figure 4A, illustrating differences in symptom severity between treatments: the infected control exhibits extensive disease symptoms, whereas the *C. sorokoniana* aqueous extract-treated plants show markedly reduced symptom development, consistent with the quantitative disease assessment. Across independent experiments, the efficacy of the *C. sorokoniana* extract ranged from 13% to 37%, showing a consistent trend toward reduced disease development compared with the infected control. However, statistical significance was achieved only at the highest efficacy level (36.7%; p ≤ 0.01, Tukey’s HSD; Figure 4B). In contrast, the commercial fungicide Rival® reduced disease severity to 74.8% (25.2% treatment efficacy), although this reduction was not statistically different from the infected control.

**Figure 4.**
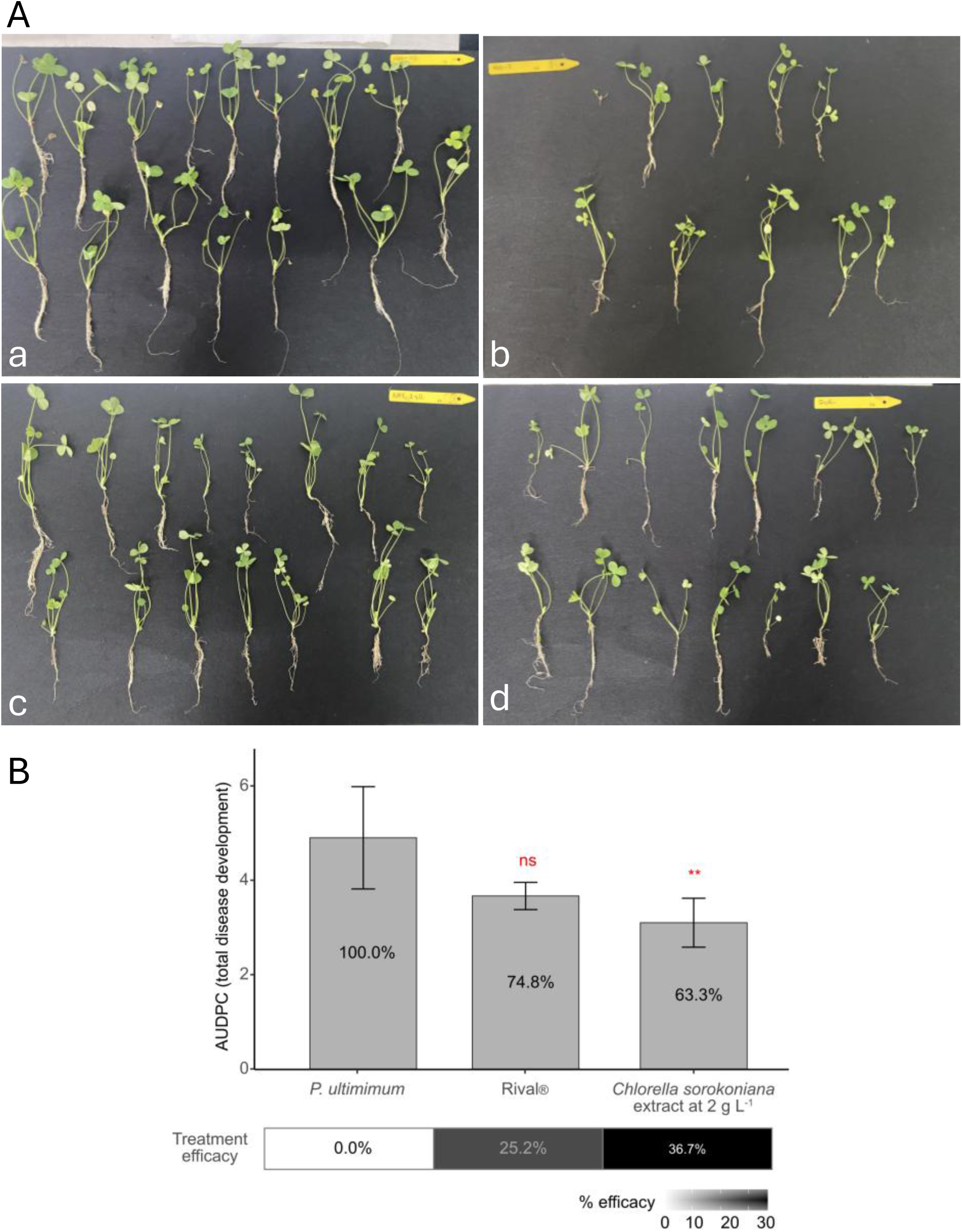
– Evaluation of the protective activity of *Chlorella sorokoniana* aqueous extract against damping-off caused by *Pythium ultimum* in Persian clover. A) Representative images of Persian clover plants at the end of the bioassay: (a) non-infected untreated control, (b) untreated plants infected with *P. ultimum*, (c) plants treated with the commercial fungicide Rival® and infected with *P. ultimum*, and d) plants treated with *C. vulgaris* aqueous extract (2 g L⁻¹) prior to infection with *P. ultimum*. B) Quantification of the disease development, expressed as the area under the disease progress curve (AUDPC), for each treatment. Values within the bars indicate disease severity relative to the untreated infected control, while the lower panel shows the corresponding treatment efficacy (%) relative to the untreated infected control. Data are presented as the mean ± SD from one experiment with five technical replicates (n = 5). Statistical comparisons were performed against the untreated infected control using one-way ANOVA followed by Dunnett’s multiple comparison test. ns, not significant; *p* < 0.01 (**).

#### 4.2.3 Cucumber-*Rhizoctonia solani* pathosystem

To further investigate the potential of *C. sorokoniana* aqueous extract to mitigate disease development by acting on the plant, we evaluated the cucumber-*R. solani* pathosystem since, as for *P. ultimum*, the extract did not demonstrate *in vitro* inhibition against *R. solani*. Representative images of the assay are shown in Figure 5A and illustrate the characteristic symptoms caused by *R. solani*, as well as the marked reduction in symptom severity following treatment with the *C. sorokoniana* aqueous extract, supporting the quantitative disease assessment. Preventive application of the aqueous extract at 1.5 g L⁻¹ reduced disease progression relative to the untreated infected control, with disease severity decreasing to approximately 45% of the control, corresponding to a treatment efficacy of 54.9% (p ≤ 0.01; Tukey’s HSD; Figure 5B). In contrast, the commercial fungicide Moncut® reduced disease severity by only 7% and did not differ significantly from the untreated infected control (Figure 5B). Across two independent experiments, treatment efficacy ranged from 48.1% to 54.9%. Although only one experiment reached statistical significance individually, both trials showed comparable reductions in disease severity, demonstrating a consistent protective effect against *R. solani*.

**Figure 5.**
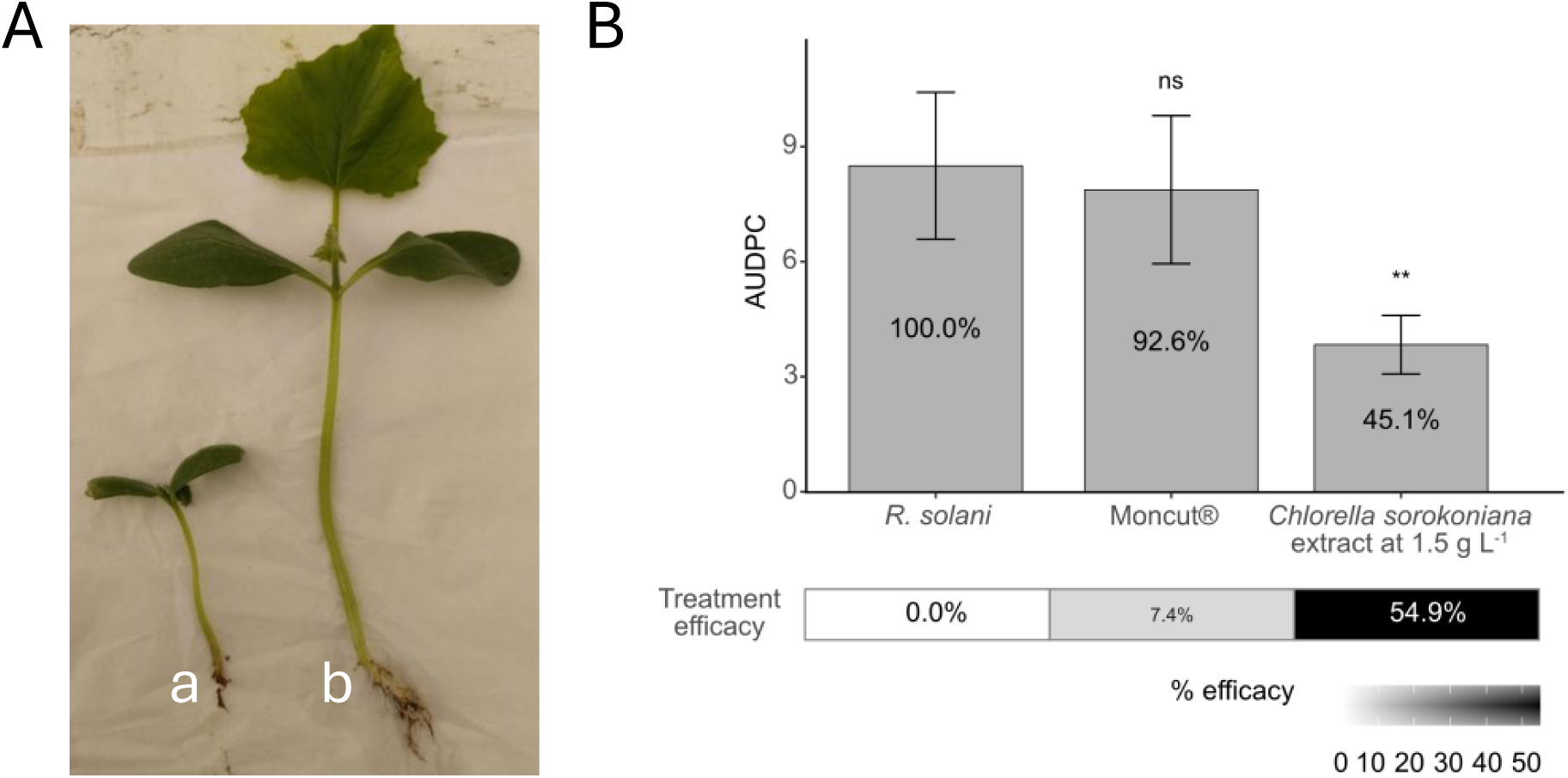
– Evaluation of the protective activity of *Chlorella sorokoniana* aqueous extract against damping-off caused by *Rhizoctonia solani* in Cucumber. A) Representative images of *R. solani*-infected cucumber plants (a) untreated and (b) treated with *C. vulgaris* aqueous extract (1.5 g L⁻¹) prior to infection. Images taken 21 days post-infection. B) Quantification of disease development in cucumber plants infected with *R. solani* and untreated, treated with the synthetic fungicide MoncutR and treated with *C. sorokoniana* aqueous extract at 1.5 g L⁻¹. Disease development is expressed as the area under the disease progress curve (AUDPC) for each treatment. Values within the bars indicate disease severity relative to the untreated infected control, while the lower panel shows the corresponding treatment efficacy (%) relative to the untreated infected control. Data are presented as the mean ± SD from two independent experiments with twenty technical replicates (n = 20). Statistical comparisons were performed against the untreated infected control using one-way ANOVA followed by Dunnett’s multiple comparison test. ns, not significant; *p* < 0.01 (**).

## 5. Discussion

### 5.1 *Chlorella sorokoniana* suppresses plant diseases through pathogen-dependent mechanisms

The present study demonstrates that an aqueous extract of *C. sorokoniana* suppresses diseases caused by pathogens with contrasting infection strategies, but through distinct biological mechanisms. While the extract directly inhibited the foliar pathogen *M. oryzae in vitro* and *in planta*, it showed little or no antimicrobial activity against the soilborne pathogens *P. ultimum* and *R. solani* at the concentrations tested *in vitro*. Nevertheless, disease development caused by both soilborne pathogens was reduced *in planta* following preventive applications. Together, these findings indicate that the extract’s biological activity is strongly dependent on the host– pathogen system and cannot be explained by a single universal mode of action. This distinction is particularly relevant because many studies evaluating microalgal products rely primarily on *in vitro* antimicrobial assays to identify promising biocontrol candidates. Our results clearly demonstrate that the absence of direct pathogen inhibition does not necessarily predict a lack of disease control under plant conditions. Similar discrepancies between laboratory screening and plant efficacy have been reported for several biological control agents (Besset-Manzoni et al., 2019), highlighting the importance of integrating both *in vitro* and *in planta* assays when evaluating natural products intended for crop protection.

### 5.2 Direct antifungal activity contributes to disease suppression in the rice-*M. oryzae* pathosystem

The results obtained for the rice-*M. oryzae* pathosystem demonstrate that direct inhibition of the pathogen is a key contributor to the disease suppression observed in both detached leaves and whole plants. The aqueous extract consistently reduced fungal biomass accumulation, impaired appressorium formation and substantially decreased rice blast severity following preventive application. Together, these complementary observations indicate that the extract interferes with critical stages of the fungal infection process, therefore limiting disease development. However, because plant responses were not investigated, the possibility that the extract also enhances host defence mechanisms cannot be excluded.

Among the observed effects, inhibition of appressorium formation is likely to play a particularly important role. Appressoria are specialised infection structures that generate the turgor pressure required to penetrate the rice cuticle and initiate host colonisation (Chakraborty et al., 2021). Consequently, disruption of appressorium differentiation is expected to substantially reduce fungal infectivity. The approximately 55% reduction in appressorium formation observed in this study is a possible mechanistic explanation for the reduction in lesion development observed in detached leaves and whole plants. The inhibition of both fungal growth and appressorium differentiation observed here extends previous reports showing that algal extracts can impair early developmental processes such as spore germination and infection establishment in phytopathogenic fungi, suggesting that microalgal metabolites may reduce pathogenicity through multiple complementary mechanisms (Bibi et al., 2026; Esserti et al., 2017).

Interestingly, the strongest antifungal activity occurred at the lowest concentrations tested (0.1– 0.2 g L⁻¹), whereas activity declined at 1.5 g L⁻¹. Although antimicrobial activity is often expected to increase with concentration, non-linear dose responses have been described for complex natural extracts, where synergistic or antagonistic interactions among bioactive metabolites can influence overall efficacy (Vaou et al., 2022). While the mechanisms underlying this response remain unknown, the reproducibility of the effect across independent assays suggests that the lower concentrations represent the optimal biological range for the antifungal activity of the *C. sorokoniana* aqueous extract.

The detached-leaf assays further support the biological relevance of the *in vitro* observations. Because infection of intact detached leaves depends on successful spore germination, appressorium differentiation and host penetration, the substantial reduction in lesion development is consistent with the inhibition of these early infection processes observed microscopically. Likewise, the high level of protection achieved following a single preventive foliar application indicates that the extract remained biologically active on the leaf surface during the initial stages of infection. Although these findings strongly support a role for direct antifungal activity, they do not exclude the possibility that the extract also primes or activates host defence responses, as has been reported for several algal-derived biostimulants and elicitors. Future studies examining defence-related gene expression and other markers of induced resistance will be required to determine the relative contribution of pathogen-directed and plant-mediated mechanisms in the *M. oryzae*–rice pathosystem.

Although the active compounds responsible for these effects remain largely unknown, *Chlorella* species produce a wide range of bioactive metabolites, including phenolic compounds, fatty acids, glycolipids, peptides, terpenoids and sulphated polysaccharides, several of which exhibit antifungal properties and/or elicit plant defence responses (Vehapi et al., 2020; Righini et al., 2022). Identification of the metabolites responsible for the observed activity will therefore be an important step towards understanding the molecular basis of the extract’s protective effects.

### 5.3 Plant-mediated responses likely contribute to protection against soilborne pathogens

In contrast to the rice blast pathosystem, disease suppression against *P. ultimum* and *R. solani* occurred despite the absence of measurable in vitro antifungal activity. This discrepancy strongly suggests that protection against these pathogens is mediated predominantly through plant-associated processes rather than direct inhibition of pathogen growth. Although the present study did not investigate these mechanisms directly, several biological processes could account for the observed protection. Microalgal extracts are increasingly recognised as effective biostimulants capable of improving root architecture, nutrient uptake, and overall plant vigour through their diverse repertoire of amino acids, polysaccharides, phytohormone-like compounds, phenolics and carotenoids (Barone et al., 2018; Farid et al., 2020; Parmar et al., 2023; Ramakrishnan et al., 2023). Enhanced root growth and plant vigour alone may increase tolerance to infection by soilborne pathogens. In parallel, numerous studies have demonstrated that algal extracts can activate plant defence signalling pathways, stimulate defence-related enzymes and induce systemic resistance against a range of pathogens (Vera et al., 2011; Prasanna et al., 2013; Roberti et al., 2015; Righini et al., 2022). Either or both mechanisms could explain why disease suppression was observed despite negligible direct inhibition of the pathogen.

Interestingly, disease reduction was consistently observed across independent experiments for both soilborne pathosystems, even though statistical significance was not achieved in every individual trial. This consistency suggests that the biological effect is reproducible but relatively moderate, as expected for plant-mediated mechanisms that rely on physiological responses in both the host and the environment rather than on direct toxicity to the pathogen. The observation of protection against two taxonomically distinct pathogens –an oomycete (*P. ultimum*) and a basidiomycete fungus (*R. solani*)– further supports the hypothesis that the extract acts primarily through host-associated processes rather than through pathogen-specific antimicrobial activity.

### 5.4 Implications for the development of microalgae-based crop protection products

From an applied perspective, the ability of a simple aqueous extract to suppress diseases caused by pathogens with contrasting lifestyles is of great interest. Unlike purified natural products, crude aqueous extracts require minimal processing, thereby reducing production costs and enhancing their potential for large-scale agricultural applications. Equally important, the present work demonstrates that efficacy should not be evaluated solely through laboratory antimicrobial assays. If only the *in vitro* experiments had been considered, the extract would likely have been discarded as ineffective against *P. ultimum* and *R. solani*, despite providing measurable protection under plant conditions. Future work should focus on identifying the bioactive metabolites, characterising the molecular mechanisms underlying plant protection, evaluating effects on the rhizosphere microbiome, and validating efficacy under greenhouse and field conditions. Formulation optimisation will also be necessary to improve stability, persistence and consistency under commercial production systems.

## Acknowledgements

The authors gratefully acknowledge Professor Nick Talbot and his laboratory (The Sainsbury Lab, Norwich, UK) for providing the Guy11 laboratory strain of *Magnaporthe oryzae*, Anne Sesma (University of East Anglia, Norwich, UK) for the pCAMBgfp vector, Pierre Gladieux and Didier Tharreau (Plant Health Institute, Montpellier, France) for providing strains isolate PR0009, and Allmicroalgae Natural Products S.A. (Leiria, Portugal) and Necton (Olhão, Portugal) for providing *C. sorokoniana* biomass.

## Funding

This work was supported by “Pacto da Bioeconomia Azul” (Project No. 16, No. C644915664-00000026) within the WP5 Algae Vertical, funded by Next Generation EU European Fund and the Portuguese Recovery and Resilience Plan (PRR), under the scope of the incentive line “Agendas for Business Innovation” through the funding scheme C5 – Capitalization and Business Innovation, by 01/C05-i02/2022.P242 to InnovPlantProtect, GreenCoLab and Fertiprado. PRR Missão Interface and FCT – Fundação para a Ciência e a Tecnologia, I.P. and Green-it Bioresources for Sustainability R&D Unit (UID/04551/2025, DOI: 10.54499/UID/04551/2025; UID/PRR/04551/2025, DOI: 10.54499/UID/PRR/04551/2025; UID/PRR2/04551/2025, DOI:10.54499/UID/PRR2/04551/2025) to InnovPlantProtect. The work of CV is supported by the Portuguese Foundation for Science and Technology for the fellowship 2024.01477.BDANA (https://doi.org/10.54499/2024.01477.BDANA). The work of LC is supported by Algarve 2030, Portugal 2030, and the European Union under the scope of AlgaeBoost—Support for the Boost and Development of GreenCoLab’s Institutional Capacity in the Area of Sustainable Blue Biotechnology, ALGARVE-FSE+-01493100. CV, LC and FG were also supported by R&D unit MED—Mediterranean Institute for Agriculture, Environment and Development (https://doi.org/10.54499/UID/05183/2025) and the Associate Laboratory CHANGE—Global Change and Sustainability Institute (https://doi.org/10.54499/LA/P/0121/2020).

## Authors Contributions

RMC coordinated the integration of research activities between GCL and InPP, the *M. oryzae*– rice studies, and the development of the data analysis strategy across pathosystems; analysed and interpreted the data; and wrote the manuscript draft. The *M. oryzae*–rice pathosystem, including the development of the GFP-expressing strain, optimisation of the infection and disease assessment protocols, disease evaluations, and statistical framework, was developed by PR, RF, CR, ACRC, and RMC. MC performed the *in vitro* assays for *M. oryzae* and *P. ultimum*, pathogen inoculum preparation, *C. sorokiniana* mode-of-action studies, and data analysis. The Persian clover–*P. ultimum* infection protocol was developed by CR, ST, and MC, with initial *in planta* evaluations performed by CR, ST and SA. The *R. solani* studies were performed by CV and BD, with CV analysing the data and contributing to manuscript preparation. Research activities were coordinated at GCL by LC, FG, and MR, and at InPP by CA and SC. CA also contributed to data interpretation and manuscript writing. All authors reviewed and approved the final manuscript.

---

**Supplementary Table 1.**
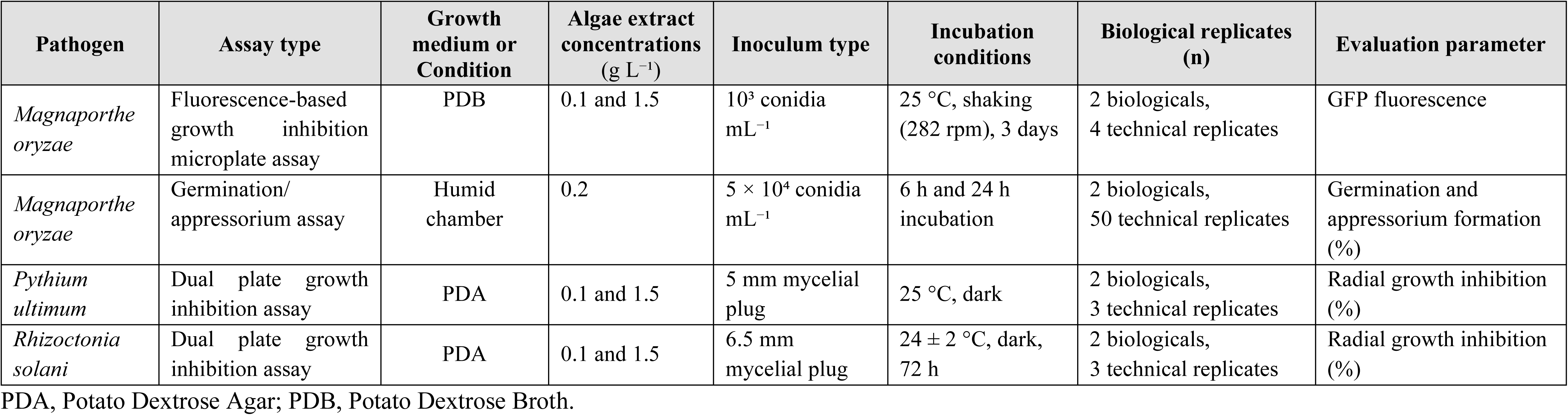
Summary of *in vitro* antifungal assays performed with *Chlorella sorokoniana* aqueous extracts against phytopathogenic fungi.

**Supplementary Table 2.**
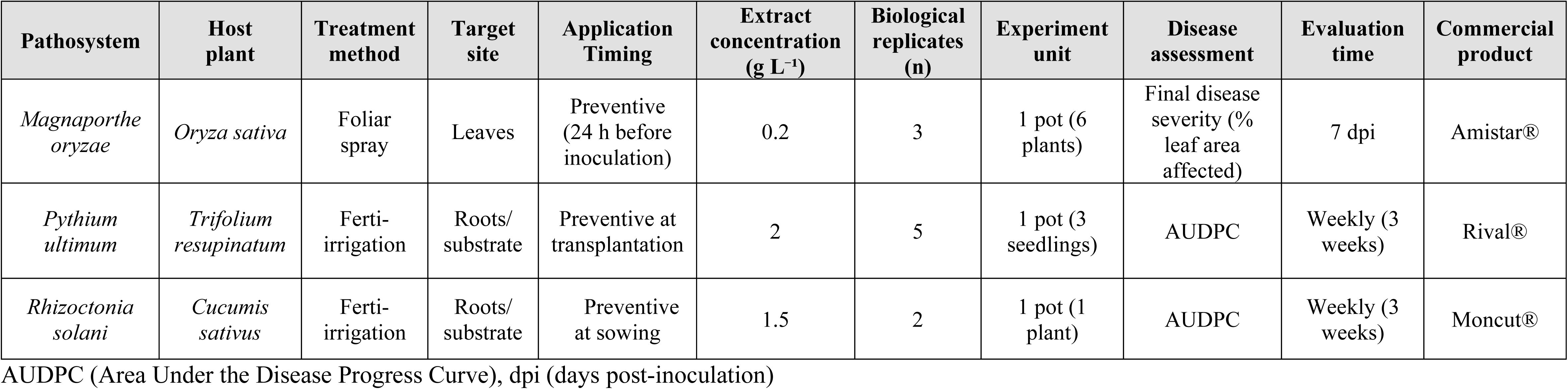
Experimental conditions and disease assessment parameters used in the *in vivo* pathosystems evaluated with *C. sorokoniana* aqueous extracts.

| Pathosystem | Host plant | Treatment method | Target site | Application Timing | Extract concentration (g L<sup>-1</sup>) | Biological replicates (n) | Experiment unit | Disease assessment | Evaluation time | Commercial product |
| --- | --- | --- | --- | --- | --- | --- | --- | --- | --- | --- |
| *Magnaporthe oryzae* | *Oryza sativa* | Foliar spray | Leaves | Preventive (24 h before inoculation) | 0.2 | 3 | 1 pot (6 plants) | Final disease severity (% leaf area affected) | 7 dpi | Amistar® |
| *Pythium ultimum* | *Trifolium resupinatum* | Ferti-irrigation | Roots/ substrate | Preventive at transplantation | 2 | 5 | 1 pot (3 seedlings) | AUDPC | Weekly (3 weeks) | Rival® |
| *Rhizoctonia solani* | *Cucumis sativus* | Ferti-irrigation | Roots/ substrate | Preventive at sowing | 1.5 | 2 | 1 pot (1 plant) | AUDPC | Weekly (3 weeks) | Moncut® |

AUDPC (Area Under the Disease Progress Curve), dpi (days post-inoculation)

**Supplementary Figure 1.**
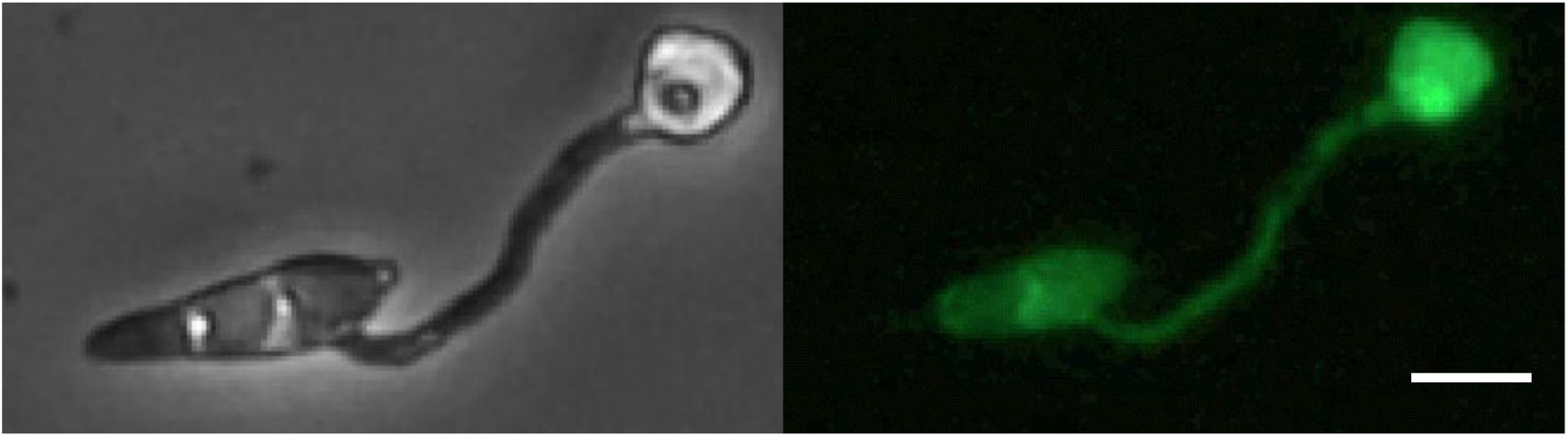
– Representative bright-field (left) and GFP fluorescence (right) images of *Magnaporthe oryzae* PR0009-GFP. Scale 5 µm.

**Supplementary Figure 2.**
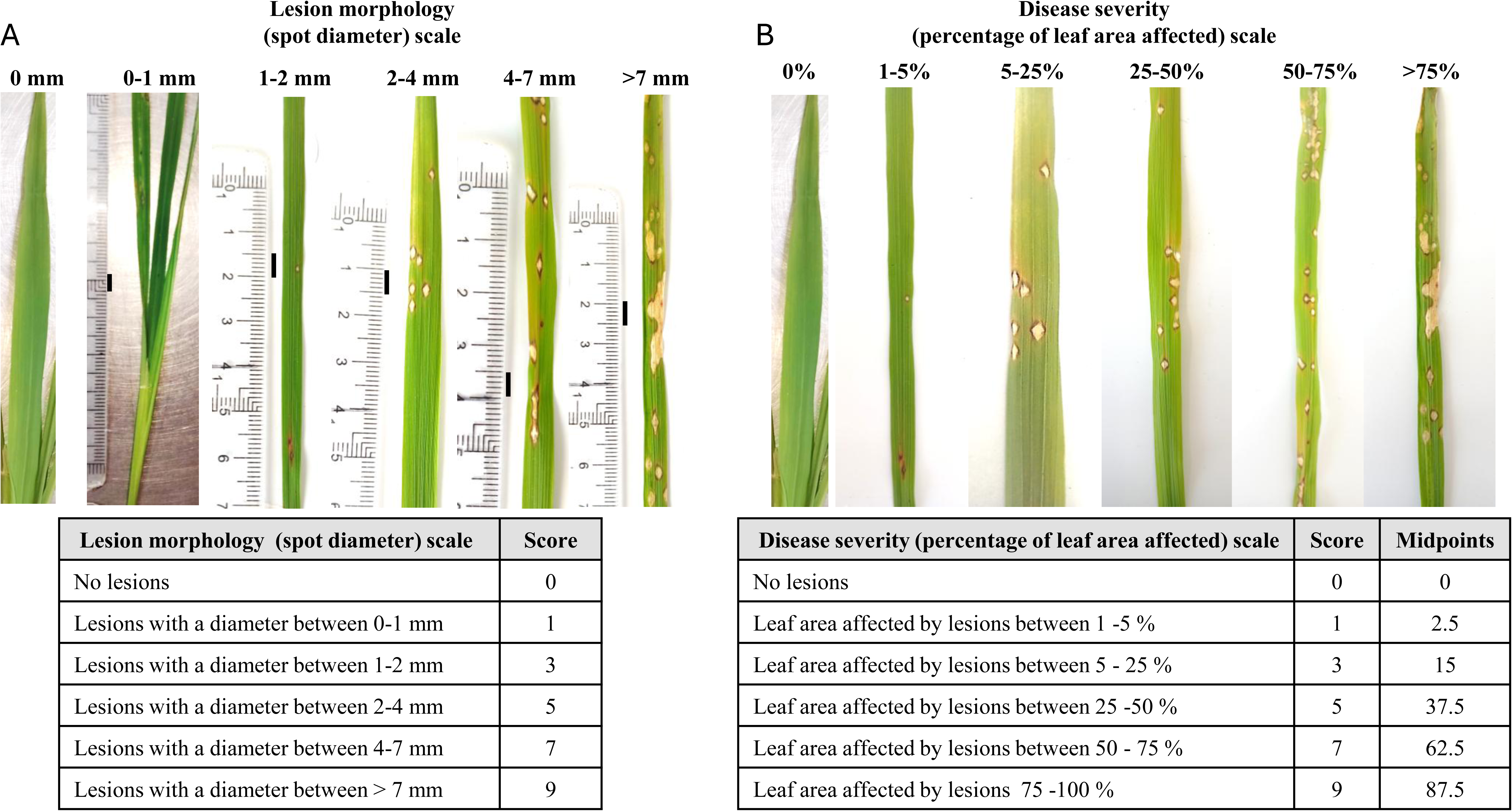
– Disease assessment developed to evaluate rice blast severity caused by *M oryzae*. Representative photos illustrate the two complementary assessment scales developed for disease evaluation: A) lesion morphology (spot diameter) showing the progression from healthy leaves to increasing lesion size and development (scale bar = 5 mm) and B) disease severity (percentage of diseased leaf area), illustrating increasing proportions of leaf area affected by blast symptoms. Representative photos correspond to the disease classes used for scoring. The spot diameter scale can be combined with the percentage-based severity scale to calculate disease indices, whereas the percentage-based scale was used for quantitative statistical analyses after conversion of ordinal scores to class midpoint values.

**Supplementary Figure 3.**
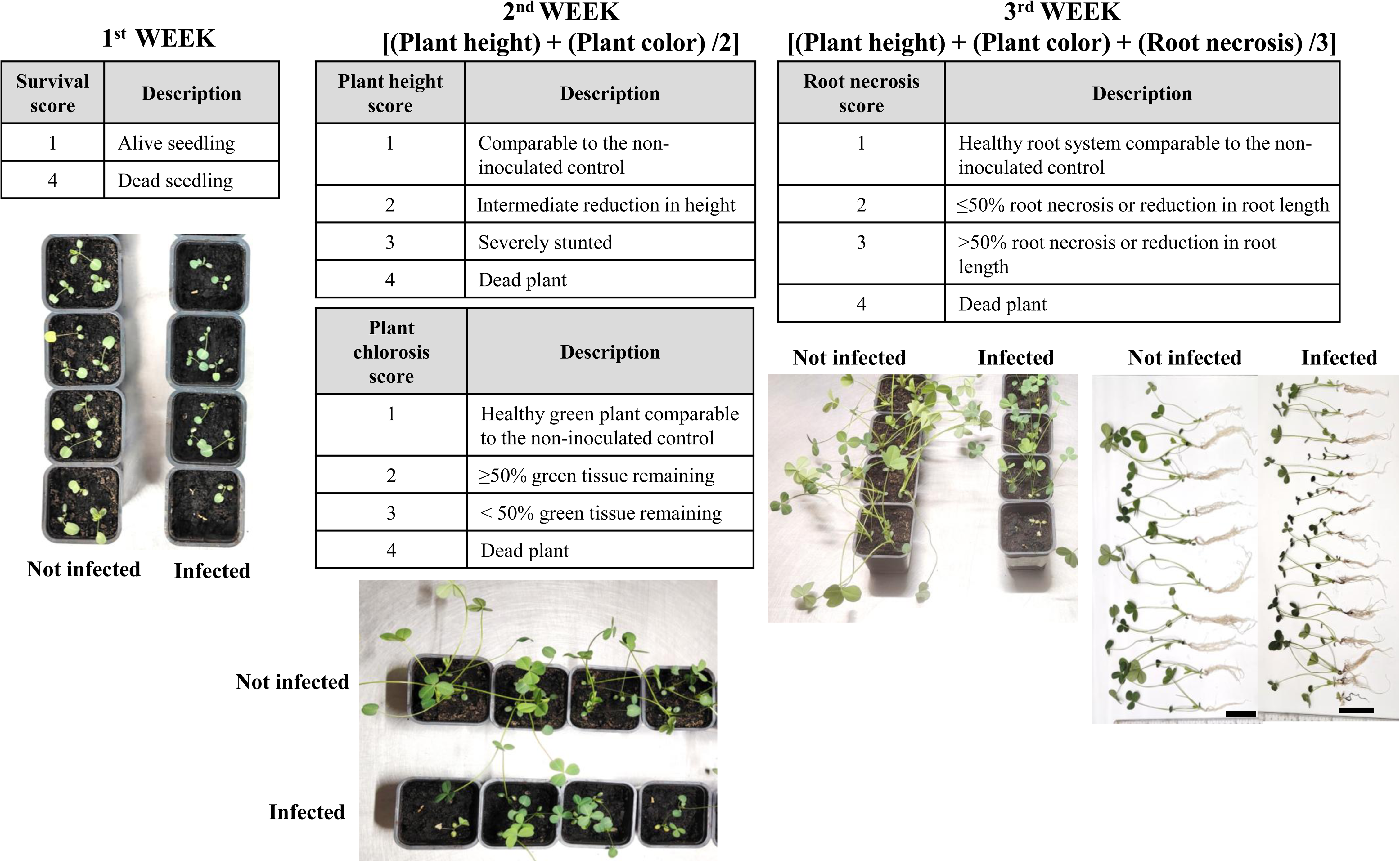
– Disease assessment developed to evaluate *Pythium ultimum* symptoms. Disease severity scale used to assess damping-off caused by *P. ultimum* in Persian clover (*Trifolium resupinatum*) over a three-week period. Representative photos show the progression of disease symptoms from seedlings to severely affected plants. Scale bar = 5 cm. Disease severity was visually assessed with a four-point ordinal severity scale based on above– and below-ground symptoms, including reduced survival, chlorosis, plant height, root necrosis, and plant death.

**Supplementary Figure 4.**
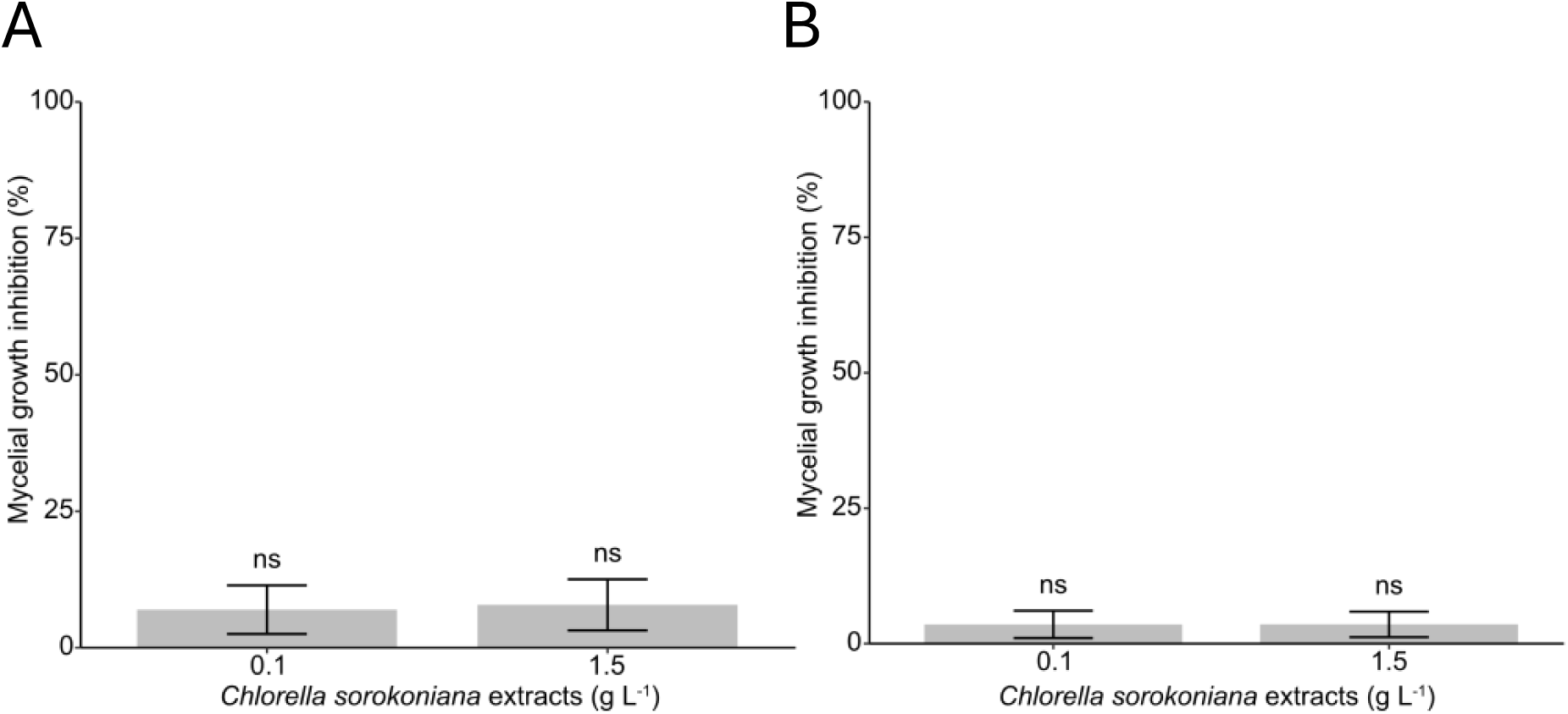
– *In vitro* effect of *Chlorella sorokoniana* aqueous extracts at 0.1 and 1.5 g L⁻¹ on mycelial growth inhibition (%) of A) *Pythium ultimum* and B) *Rhizoctonia solani*. Data represent the pooled results of two independent experiments (three technical replicates per experiment) and are presented as the mean ± SD. Each treatment was compared with the untreated control using one-way ANOVA followed by Dunnett’s multiple comparison test. ns, not significantly different from the untreated control (*p* > 0.05).

## Notes

### Competing Interest Statement

The authors have declared no competing interest.

